# Echosounders for fish detection disturb harbour porpoises

**DOI:** 10.1101/2025.11.03.686290

**Authors:** Jeroen Hubert, Benoît Bergès, Clea Parcerisas, Elisabeth Debusschere, Joost A. C. de Bruijn, Jozefien M. Demuynck, Carlota Muñiz, Hans Slabbekoorn

## Abstract

The marine world is an acoustic world that has become noisier with the increasing diversity and intensity of human activities at sea. The harbour porpoise (*Phocoena phocoena*) is one of the best-studied cetaceans regarding hearing and responses to human-made underwater sound. The species is most sensitive to high frequencies, yet most impact studies have focused on relatively low-frequency sources. As such, the effects of high frequency sonar – including echosounders – remain largely unstudied, despite their widespread use on vessels for depth sounding, fish or object detection, and seabed mapping. We investigated the effects of scientific echosounder use on harbour porpoise occurrence using 13 multi-sensor mooring deployments in the southern part of the North Sea. Moorings operated for an average of 57 days, with split-beam scientific echosounders active for an average of 51 days, for 10 minutes every hour (5 min at 70 kHz, followed by 5 min at either 185–255 kHz or 70 kHz again). Porpoise acoustic presence was continuously monitored with C/F-PODs and additionally validated with hydrophone detections at four of the locations. Across all 13 sites, detections declined by 65– 79% during echosounder transmissions and returned to typical levels within ∼30 minutes after the echosounder stopped pinging. Despite this relatively quick recovery, there was no indication of habituation, as responses did not diminish across observation periods over six weeks of hourly exposure. Spatial effects appeared local, as no deterrent effect was observed at 2.5 km from the source. We believe that our findings have important implications for studies that investigate both harbour porpoise and fish presence to understand predator-prey interactions, but they should also raise concern about the potentially cumulative impact on sensitive cetaceans from the wide use of relatively high frequency sonar in offshore practices.

## Introduction

Marine animals as well as humans use underwater sound to orientate, communicate, and detect food and threats in the marine environment (Ladich, 2019; Lurton, 2010). Passively, by just listening, and actively by echolocation in animals and through active sonar systems by humans (Erbe et al., 2025b; Wisniewska et al., 2015). Such use of sound allows animals and humans to obtain information about their surroundings, even in the absence of light, at depth, or at night. Humans therefore share – and sometimes compete for – the same acoustic space as marine animals, adding to the already noisy soundscape from a wide range of human activities at sea (Duarte et al., 2021; Slabbekoorn et al., 2010). Understanding how these sounds affect marine life has become an important focus of acoustic research, using both passive and active acoustics (Kok et al., 2021; Watson et al., 2025).

Prominent human-made noise sources include shipping, seismic surveys, piling, drilling, and explosions. These sources produce most of their acoustic energy in the lower frequencies, which can be detected by virtually all marine animals including invertebrates and fish (Duarte et al., 2021). Such noise has been shown to disrupt communication (Radford et al., 2014), orientation (Lecchini et al., 2018), foraging (Hubert et al., 2018), and cause stress (Debusschere et al., 2016) and displacement (van der Knaap et al., 2021). Marine mammals, and especially toothed whales, can also hear much higher frequencies due to their ability to echolocate (Southall et al., 2019). This sensitivity is exploited by acoustic deterrent devices such as high-frequency pingers, which are used by fishermen to mitigate dolphin bycatch and depredation (Bruno et al., 2021; Buscaino et al., 2021), though concerns about excessive displacement and hearing impairment exist (Dolman et al., 2022). Ultrasonic antifouling devices have also been shown to deter Cuvier’s beaked whales (*Ziphius cavirostris*) (Trickey et al., 2022). So, the wide hearing range of marine mammals makes them vulnerable to different anthropogenic sound sources, including sonar systems.

Active sonar systems are widely used for object detection and mapping in the underwater environment. Sonars generate series of sound pulses and analyse the returning echoes to detect targets such as submarines, seafloor features, or fish schools. Military sonars often operate at low- or mid-frequencies (<2–3 and <8–20 kHz, resp.), whereas other sonar systems including echosounders are primarily used for scientific and commercial applications and operate at higher frequencies (>10 – >1.000 kHz) (Erbe et al., 2025b). Military sonars typically emit long signals (up to several seconds) that propagate over large distances, whereas higher frequency systems typically produce short (typically less than a few milliseconds) and highly directional pulses with a much shorter range, and often at high repetition rates. The use of military sonar has been associated with mass stranding events of cetaceans, particularly beaked whales, although direct cause-effect relations are often difficult to establish (Evans et al., 2001; Parsons et al., 2008). Military sonar exposure can also result in (temporary) hearing loss (Finneran, 2015) and, at more subtle levels, can trigger physiological and behavioural responses in cetaceans (Harris et al., 2018; Pirotta et al., 2021). The potential impact of echosounders received less attention, despite its common use.

The impact of echosounders has been studied in a few toothed whale species (Erbe et al., 2025a). Short-finned pilot whales did not leave the area or stop foraging during 38 kHz echosounder activity, but they did alter their heading more frequently (Quick et al., 2017). For four species of beaked whales (grouped together), acoustic – but not visual – presence decreased during exposure to a multi-frequency echosounder (18, 38, 70, 120, and 200 kHz), potentially explained by reduced foraging behaviour (Cholewiak et al., 2017). Similarly, the acoustic presence of Cuvier’s beaked whales was lower during detections of echosounder use (mainly 28 and 50 kHz) (Trickey et al., 2022). In contrast, another study on the same species found no impact of 12 kHz multibeam echosounder surveys on their foraging behaviour (Varghese et al., 2021). Pilot and beaked whales are high-frequency hearing cetaceans, highlighting the need to investigate the effects of echosounders on species with even higher frequency hearing, such as harbour porpoises (Southall et al., 2019).

Harbour porpoises are widely distributed in temperate coastal waters of the Northern hemisphere, where they play an important ecological role as mesopredator (Das et al., 2003). They are a sensitive, very high-frequency hearing cetacean – hearing up to at least 180 kHz – (Kastelein et al., 2002; Southall et al., 2019) and their responses to human-made noise are well-studied (Erbe et al., 2025a). They show a high seasonality in the Southern Bight of the North Sea, with high presence during winter season and low presence in summer (Calonge et al., 2024). Their relatively small size in cold waters necessitates a high metabolic rate, requiring near-continuous foraging (Wisniewska et al., 2016). Because they heavily rely on high-frequency echolocation clicks for navigation, foraging, and communication, they are particularly vulnerable to acoustic disturbance (Sørensen et al., 2018a; Wisniewska et al., 2016). Widely used echosounders may therefore be an important but understudied source of disturbance. Understanding whether echosounders influence porpoise vocal activity is highly relevant, as reductions in acoustic presence could indicate avoidance behaviour, reduced foraging, or altered social interactions (Bergès et al., 2019). Despite their prevalence, the behavioural effects of echosounders on harbour porpoises remain poorly understood. Addressing this gap is essential for advancing scientific knowledge of human-wildlife interactions and for informing management practices aimed at balancing fisheries and conservation.

In the current study, we investigated the effect of bottom-moored split-beam echosounders, deployed to monitor presence and abundance of pelagic fish, on the vocalization of harbour porpoises. We quantified porpoise acoustic presence during periods with and without echosounder transmissions using 13 long-term deployments of multi-sensor moorings equipped with echosounders, acoustic cetacean loggers (C/F-PODs), and hydrophones. In addition, we used data from two C-PODs from another project deployed at ∼2.5 km from our moorings to check for longer-range impact. The replicated nature of multi-week exposure data and sufficient data with and without echosounder operation allowed us to answer the following questions: Does echosounder activity reduce the acoustic presence of harbour porpoises? Is such a reduction in acoustic presence caused by reduced call activity or leaving the area? Is there a significant recovery within the 50-minute period in between subsequent echosounder exposure events? Does this recovery reach baseline levels before the next echosounder activity? Is there a different impact of an echosounder with a 70 kHz signal, from an echosounder with a 185-255 kHz signal? And is there any sign of habituation over the weeks of hourly exposure events?

## Materials & Methods

### Study sites

This study is based on data originally collected to investigate the effects of offshore wind farms on pelagic fish and harbour porpoises (project: APELAFICO). We used data from 13 long-term deployments of multi-sensor moorings in the Southern Bight of the North Sea. Deployments took place in offshore wind farms and near shipwrecks, both located in Belgium and the Netherlands (Figure 1, Table A1). Moorings were deployed for an average duration of 57 days (range: 40–71 days), in (3 ×) summer 2021, and in (4 ×) spring, (4 ×) summer, and (2 ×) autumn of 2023 (Figure 1).

**Figure 1:**
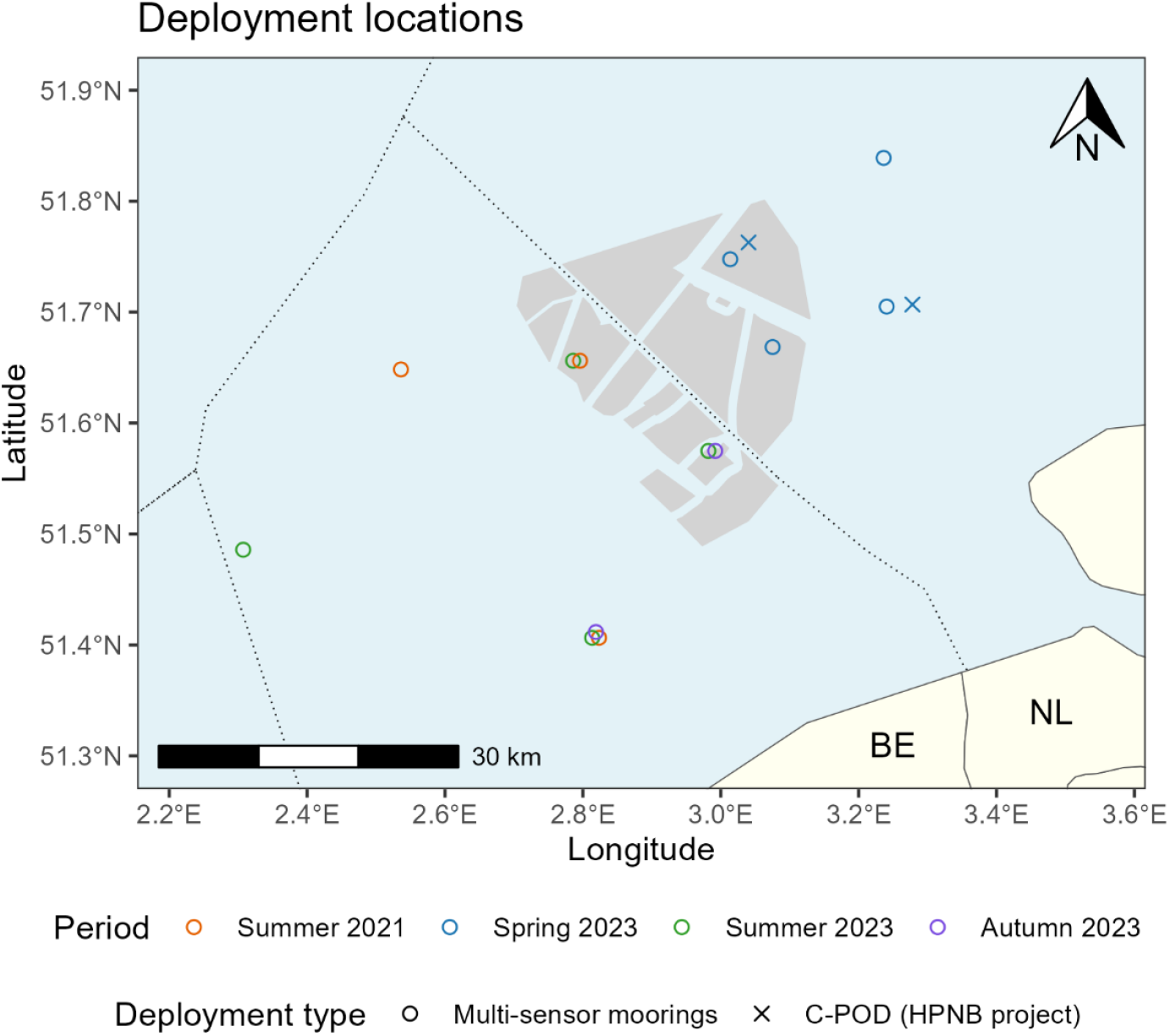
Map of the study area off the coasts of Belgium (BE) and the Netherlands (NL). 13 multi-sensor mooring deployments of the APELAFICO project are indicated by open circles, and two selected locations of the HPNB project by cross marks, with the colours indicating the deployment periods. Identical deployment positions are shown as slightly offset to ensure visibility of all deployments. Grey polygons represent offshore wind farms and grey dotted lines indicate the Exclusive Economic Zones.

Our data collection in spring 2023 in the Netherlands overlapped with a long-term monitoring campaign on harbour porpoise habitat use in this area, for a project called Harbour Porpoise Network Borssele (HPNB; Olivierse et al. 2024). We used data from two stations from this campaign for a modest test of the spatial range of the observed effects. The locations of these stations were at 2.48 and 2.58 km from two of our moorings (Figure 1).

### Multi-sensor moorings

We used four bottom-mounted multi-sensor moorings (1.6 × 1.2 × 1.0 m; L × W × H), all equipped with an echosounder, cetacean logger, hydrophone, and acoustic release system.

The scientific echosounders (Wide Band Autonomous Transceiver - WBAT, Kongsberg Maritime AS, Bergen, Norway) were originally used to monitor pelagic fish abundance. However, in the context of this study, the echosounders were regarded as treatment rather than sensors. Each echosounder was equipped with an upward-pointing split-beam transducer with an 18° angle beamwidth and set to ping in continuous wave mode at 150 W with a nominal frequency of 70 kHz (ES70-18CD, Simrad, Horten, Norway), and a split-beam transducer with a beamwidth of 7° set to produce frequency modulated (upsweep) broadband pulses at 75 W in the 185 to 255 kHz range (ES200-7CDK-split, Simrad). A complete overview of echosounder settings is provided in Table A2. Echosounders were used for an average period of 51 consecutive days (range: 40 – 60 days) during deployments. During this period, echosounders operated for 10 minutes at the start of every hour: In 11 out of 13 deployments, we used the ES70 transducer in the first five minutes, followed by the ES200 transducer in the second five minutes; In two out of 13 deployments, only the ES70 transducer was used, for the first 10 minutes of each hour (Figure 2a).

**Figure 2:**
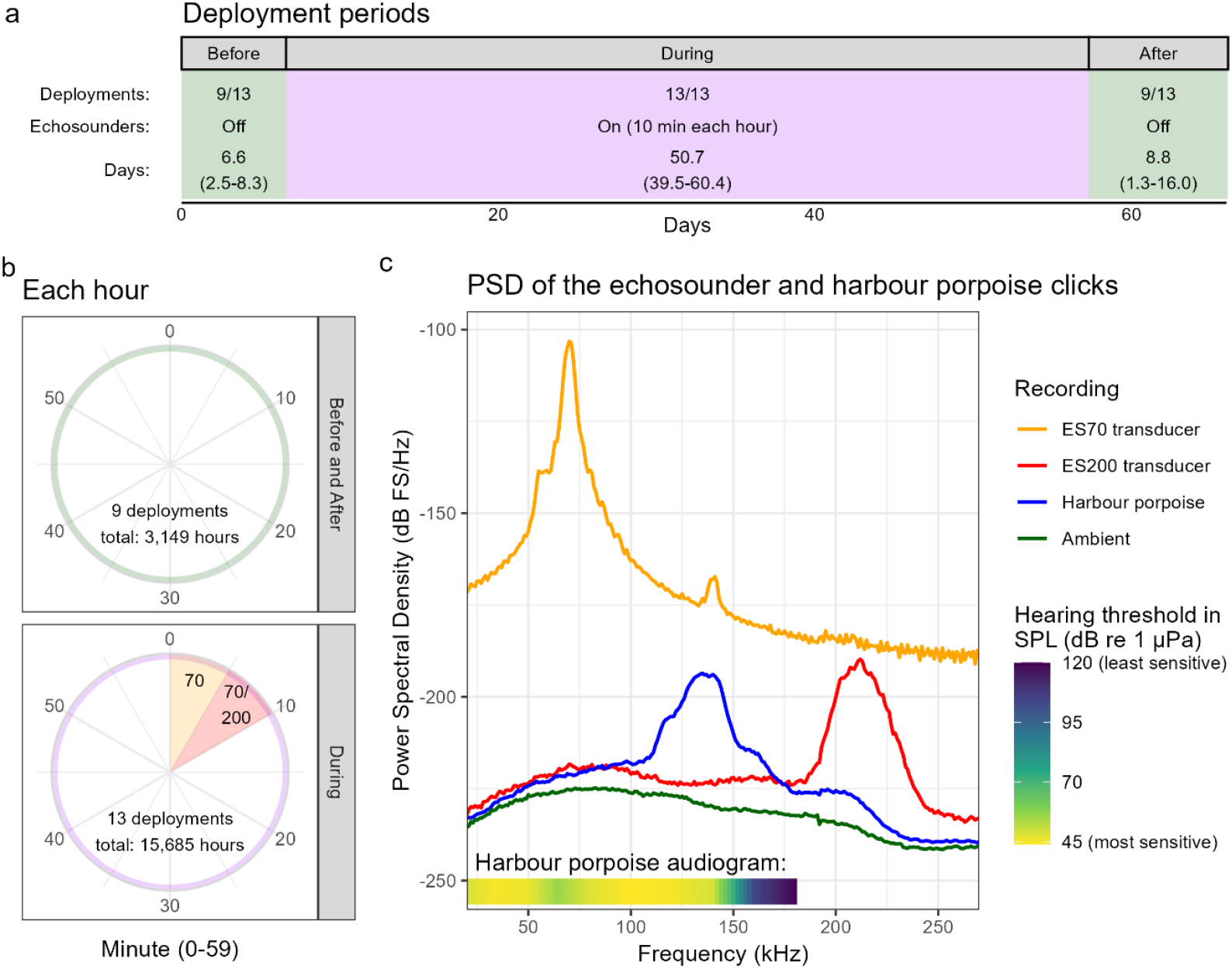
(a) Overview of deployments. Each deployment had a single During period in which the echosounder was used for 10 minutes each hour, and a Before and After period without echosounder activity, which were merged for the analyses. In 4 out of 13 deployments, the combined Before and After period was deemed too short (<1 day) to be included. The bottom row ‘Days:’ shows the mean number of days and minimum and maximum for all included deployments. The C/F-PODs were active across all periods. (b) Overview of each hour in the Before and After period, and the During period. In the During period, each hour started with 10 minutes of echosounder use, always first 5 minutes with the ES70 transducer followed by another 5 minutes of ES70 or 5 minutes of ES200. (c) Relative power spectral density (PSD) plot for various sound types recorded by the SoundTrap click detector. Coloured lines indicate the log-energy mean of a sound type for all four deployments with high click detection. Note that one out of four deployments recorded at a lower sample rate (384 vs. 576 kHz), hence the ES200 transducer could only be recorded in one deployment. The absolute differences in amplitude may be explained by differences in distance between transducer and SoundTrap. The bar at the bottom shows the hearing sensitivity of the harbour porpoise (from Kastelein et al. 2002; Kastelein et al. 2010) in the 20-180 kHz range (their actual audiogram starts at 250 Hz, not included here).

The cetacean loggers (C-PODs in 12 deployments, and an F-POD once, Chelonia Inc., Cornwall, UK) recorded the acoustic presence of odontocetes continuously. However, the C-PODs (and not the single F-POD) was set to record a maximum of 4096 high-frequency clicks per minute. The signal from the ES70 transducer was also detected by the filter settings and saturated this capacity, resulting in approximately ∼49-55 minutes of usable data per hour (the full minutes without activity from the ES70 transducer).

The hydrophones were various SoundTrap models (Ocean Instruments, Auckland, New Zealand), either a ST300 HF, ST600 HF, or ST4300 HF with four external HTI-96-MIN hydrophones (High Tech Inc., Long Beach, MS, US). These recorded ambient sound at a 48.000 Hz sampling rate, for 10 to 60 minutes per hour. Ambient recordings were not used in this study, but when the hydrophone model allowed (300 & 600 HF), the SoundTrap Click Detector function was also enabled to record potential clicks of odontocetes. When this function was enabled, and these hydrophones were recording for 60 min per hour, the detections were used for this study; which was the case for four deployments (Table A1).

Lastly, an acoustic release system (VR2AR, VEMCO, Billings, MT, US) enabled the retrieval of each mooring at the end of the deployment.

### HPBN stations

The two selected HBNB stations consisted of a continuous recording C-POD, attached to a rope between an anchor and sub-surface buoy.

### Echosounder signal characterization

To characterize the echosounder signal and put it into context, we determined the spectral content of high click detections by the SoundTraps from four deployments (Table A1). For each deployment and sound type (harbour porpoise, echosounder transducers ES70 and ES200, and ‘other sound’); up to 10,000 snippets were randomly selected when available. For each snippet, the PSD was computed using the signal.periodogram function from the spicy library in Python (Virtanen et al., 2020), with a Kaiser window and an FFT length set to one-thousandth of the sampling rate. PSD values are given in relative values, as the calibration response by the manufacturer is only valid up to 60 kHz.

The resulting power spectral density plot (Figure 2b) shows that the spectral energy of the ES70 transducer (with current settings, see Table A2) peaks at 70 kHz with a minor harmonic tone at 140 kHz. The signal of the ES200 transducer concentrates between 192 and 233 kHz, well above the range of sensitive hearing for harbour porpoises (Kastelein et al., 2010, Kastelein et al., 2002). Harbour porpoise clicks concentrate in the 114-150 kHz range. ‘Other sound’ typically matched sound types outside the frequency range of their expected signal, suggesting the majority of these snippets can be interpreted as (close to) ambient levels. Note that absolute differences in levels should be interpreted with care as SoundTraps were positioned at varying distances from the transducers (0.1 – 0.5 m). They were also at the same height on the moorings as the transducers, while most acoustic energy of the echosounder – especially from the ES200 transducer – went upwards (Simrad, n.d., n.d.).

### Porpoise detections processing

We processed the harbour porpoise detections by the C/F-PODs and SoundTraps to gain insight into the acoustic presence and behaviour of harbour porpoises in relation to the active echosounder periods. For the C/F-PODs, click trains were first classified by the C/F-CPOD software using the KERNO classifier (Chelonia Limited, 2014). All click train qualities were included. Individual clicks were further classified into regular clicks (regular clicking for navigation or prey searching), buzz clicks (prey capture or social communication, Miller 2010; Clausen et al. 2011; Sørensen et al. 2018) and inter-train intervals (i.e. pauses between click trains) based on the inter-click-interval length using a three components Gaussian mixture model (Bergès et al., 2019).

For the SoundTraps, the output from the HF Click Detector detects potential clicks by using a simple energy thresholding in the frequency band between 115 kHz and 160 kHz. Each of these clicks was then a-posteriori classified into noise, low-, or high-quality porpoise click using the model developed and described in Cosentino et al. (2019), by means of the Python package pyporcc (Parcerisas, 2022). Only clicks classified as high-quality porpoise clicks were included in the analysis. As SoundTrap click detections were only available for four out of 13 of the multi-sensor mooring deployments, the main goal of this analysis was to verify the patterns found in the C/F-POD data.

Further data processing was conducted in R (R Core Team, 2025). We used the echosounder (‘sonar’) detections by the C/F-PODs to correct for clock-drift between the echosounder and C/F-PODs. This correction step was not applied to SoundTrap click detections, as these were used as a qualitative validation of the C/F-POD data only. Subsequently, we calculated the proportion of minutes with harbour porpoise click detections for each minute of the hour, by aggregating across all hours per deployment. This metric is referred to as the proportion of porpoise positive minutes (PPM). We calculated the proportion PPM separately for each deployment, and for two periods: (1) the combined period Before and After echosounder activity, and (2) the period During echosounder activity. This procedure was applied to both C/F-POD detections (both from our multi-sensor mooring deployments and from the HPNB deployments), and SoundTrap detections (from our multi-sensor moorings).

Besides the proportion PPM, we also calculated the mean number of detected clicks per positive minute and examined trends in the same way as proportion PPM to gain insights into whether trends in porpoise detections could be explained by changes in their click rate rather than their physical presence. To assess whether harbour porpoise occurrence returned to typical levels during the 50-minute pauses in between echosounder activity, we compared the mean proportion PPM in the last 20 minutes of these pauses (in the final week of echosounder operations) with the mean proportion PPM recorded 3 to 7 days (depending on data availability) after the echosounder activity had ceased. These relatively short periods of a week were chosen to minimize the influence of seasonal patterns. To gain insight into potential changes in echosounder impact over time in the During period, we calculated the proportion PPM per week since the start of the During period; a weighted average of all deployments was computed for each week. To account for variation in harbour porpoise abundance between weeks, we normalized the proportion PPM by dividing each minute’s proportion PPM by the mean proportion PPM of that week. Lastly, to explore potential changes in porpoise behaviour, we examined the proportion of click type classifications (‘buzz’ or ‘other’) over time and between periods. We plotted and analysed this in 5-minute bins rather than per minute to prevent that very low numbers of click detections would skew the data.

Due to saturation in click detections by the C-PODs when the ES70 echosounder transducer was active, we excluded data from these periods for the statistical analyses. A one-minute safety margin was added (i.e., data from minute 59 up to and including minute 5 or 10) to account for potential – though likely minimal – errors in clock drift correction.

### Statistics

We fitted generalized additive models (GAM) with quasibinomial error distribution and logit link function for the proportion PPM and proportion buzz clicks, and with a Gamma distribution and inverse link functions for click counts. To investigate how these response variables varied within the hour, we used minute of the hour as a smooth term predictor. Depending on the specific research question, we additionally included a categorical variable: the period (Before and After vs. During, Figure 2a), echosounder configuration (ES70 + ES200 vs. ES70 only), or the number of weeks since the echosounder start. Deployment ID was always added as random effect using a random smooth.

We compared model structures by evaluating whether fixed effects improved model fit, either as additive terms or as interactions with the smooth term for minute of the hour. Models were compared using analysis of deviance tables, and the best-fitting model was selected based on lowest residual deviance and statistical support over simpler models. The basis dimension (k) of smooth terms was evaluated post hoc to ensure it was not set too low relative to the effective degrees of freedom.

From the selected models, we reported parametric coefficients, estimated degrees of freedom (edf), F-statistics, p-values for smooth terms, deviance explained, and adjusted R^2^ values. When the smooth term for minute of the hour was statistically significant, we plotted its estimated partial effect for each category level, and we assessed at which minutes the differences between categories were significant. These comparisons were visualized using confidence intervals on the pairwise differences between smooths in the appendix.

For the assessment on whether harbour porpoise occurrence returned to typical levels during the 50-minute pauses between echosounder activity (Figure 4), we used a paired t-test comparing the proportion PPM between two time-bins. Prior to analysis, we verified that the differences between paired observations were normally distributed. We reported the t-value, p-value, and mean difference with 95% confidence interval.

## Results

Harbour porpoise acoustic presence was lowest during and immediately after the 10-minute echosounder activity, gradually increasing and stabilizing during the 50-minute period without echosounder signal before dropping again when echosounders were reactivated (Figure 3a). No consistent within-hour patterns were observed during days without echosounder activity (Figure 3b). Across all 13 deployments, the weighted mean proportion of porpoise positive minutes (PPM) in the second minute following echosounder activity was 0.013, rising to 0.036 in the final 20 minutes before the next echosounder activity: an increase of 182%, followed by a 65% drop when the echosounder restarted. This pattern was even more pronounced when focusing on the second minute after the ES70 transducer, with a mean proportion PPM as low as 0.008 which means a 370% rise and then fall of 79% with respect to the final 20 minutes of the hour (Table A3).

**Figure 3:**
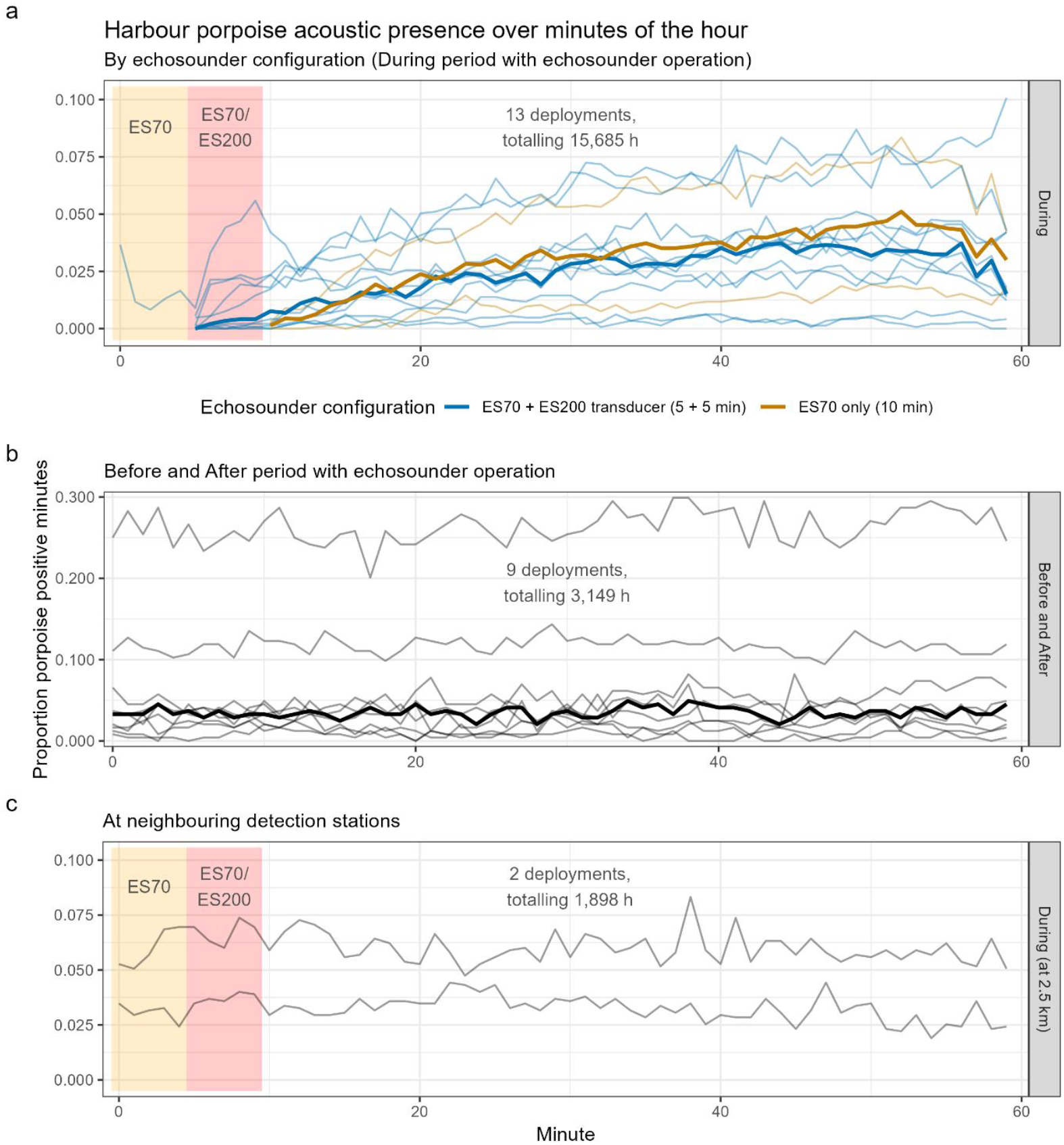
Proportion of harbour porpoise positive minutes (PPM), for each minute of the hour. The thin lines indicate the individual deployments, and the thick lines indicate the medians of all locations together. The yellow and red shaded areas indicate the minutes with echosounder activity: first 5 minutes by the ES70 transducer, followed by 5 minutes of the ES200 transducer (in 11 deployments), or another 5 minutes of ES70 (in two deployments). (a) In the During period with echosounder operation, acoustic presence of harbour porpoises is lowest during and right after echosounder operation and then recovers to stable levels in the last ∼20 minutes of the hour. The colour of the lines indicate the echosounder configuration. The recovery after ES70 only lagged behind slightly yet significantly, but quickly caught up with recovery after ES70 + ES200. This panel is based on 13 deployments totalling 15,685 h of data. During minutes with an active ES70 transducer, proportion PPM data are an underestimation because of C-POD (but not F-POD) saturation by the echosounder. (b) Before and After the period with echosounder operation, the acoustic presence of harbour porpoises is stable within the hour. This panel is based on 9 out of 13 deployments, totalling 3,149 h of data. (c) At ∼2.5 km distance from our echosounder, also no within-hour pattern in harbour porpoise acoustic presence was found at two stations from the HPNB project, totalling 1,898 h of data. Note that the y-axes differ in scale.

**Figure 4:**
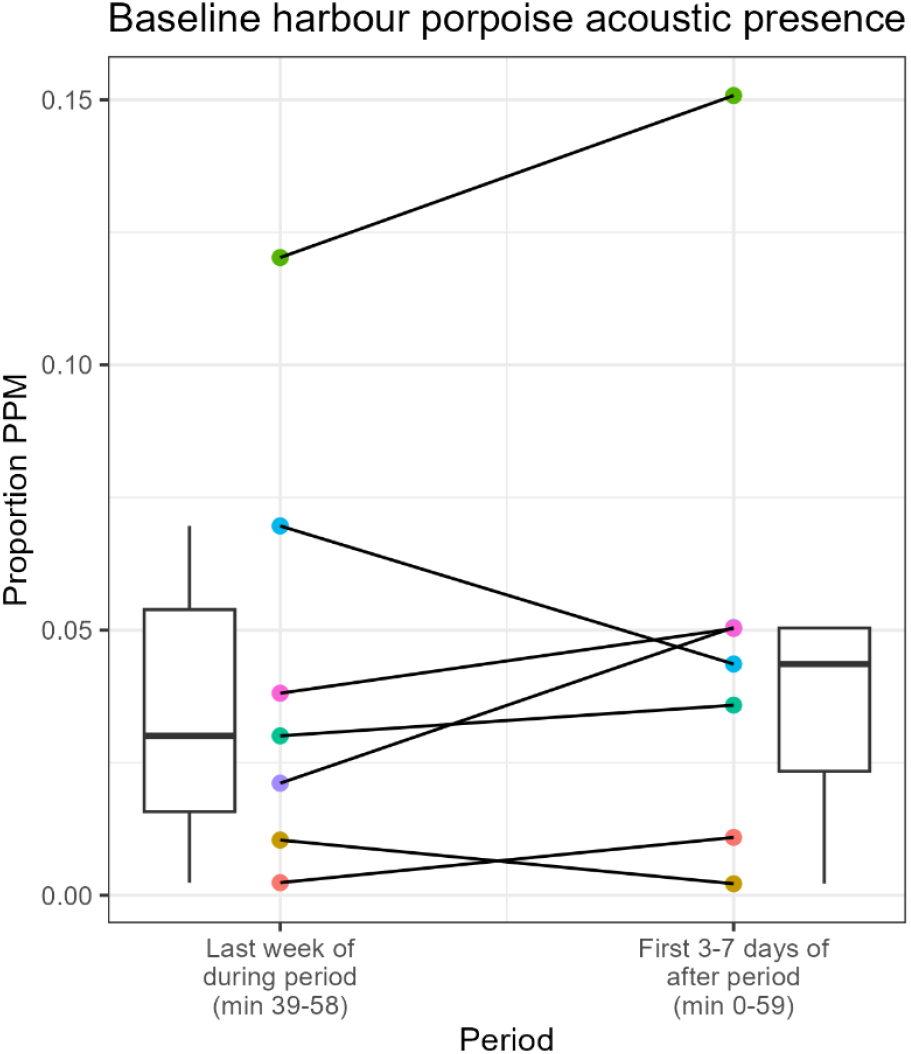
Comparison between the mean proportion PPM of min 39-58 in the last week of echosounder activity (the During period) and the mean proportion PPM of min 0-59 in the first 3-7 days of the after period. No differences between periods were found, indicating that the acoustic presence of harbour porpoises between echosounder activity recovered to typical levels. The coloured datapoints are the means from seven different deployments; only deployments with three or more days after the echosounder activity are included, when possible, we used the first seven days to compute the means after echosounder activity.

This pattern was statistically confirmed by a generalized additive model (GAM) that showed a significant non-linear effect of minute of the hour on the proportion PPM in the period with echosounder activity, across echosounder configurations (During: edf = 3.49, F = 42.24, p < 0.001, Figure 3a). Model estimates revealed that porpoise acoustic presence first increased and then stabilized in the final ∼20 minutes of the hour (Figure A1). In contrast, no significant effect of minute of the hour was found in the period without echosounder activity (Before and After: edf = 1.00, F = 2.43, p = 0.119, Figure 3b & A1). Irrespective of time, there was also an overall difference between the periods, with the Before and After period having a higher proportion PPM (estimate = −0.83, ± 0.03, p < 0.001). The deployment, as random effect, was also highly significant (edf = 11.88, F = 231.93, p < 0.001), indicating substantial variation among deployments. The model explained 77.5% of the deviance (adjusted R^2^ = 0.766).

Next, we compared the impacts of echosounder transducer configuration (ES70 + ES200 vs. ES70 only). Both deployment types showed significant non-linear variation in proportion PPM over minutes of the hour (ES70 + ES200: edf = 5.47, F = 147.5, p < 0.001; ES70 only: edf = 5.92, F = 34.8, p < 0.001, Figure 3a and A5). Model estimates indicated that the increase in porpoise acoustic presence lagged behind slightly but significantly in ES70 only-deployments at first, but quickly caught up (Figure A2). The random effect for deployment was also significant (edf = 11.97, F = 352.8, p < 0.001). The model explained 91.8% of the deviance (adjusted R^2^ = 0.935). Given that only two deployments used ES70 only, we minimized overfitting by restricting the basis dimension (k = 12) for the smooth terms, which was sufficient given the effective degrees of freedom and model diagnostics.

The proportion PPM at two nearby C-PODs, located 2.5 km from the echosounders, did not indicate any deterrent effect at this distance, based on visual inspection of the data (Figure 3c).

These results are based on harbour porpoise detections recorded by C/F-PODs, available from all 13 deployments. For four of these deployments, harbour porpoise clicks were also detected using SoundTraps, and we processed their data similarly as cross-device validation. Visual comparison of the detection patterns over time shows that SoundTrap detections followed the very same trends as the C/F-POD detections (Figure A2). Further analyses are based on C/F-POD data as these were available for all deployments.

To explore whether the drop in acoustic detections of porpoises could be explained by a drop in their click rate, rather than a deterrence effect, we analysed the number of clicks per positive minute. This varied significantly over the hour in the During period with echosounder use (edf = 6.69, F = 8.33, p < 0.001, Figure A3 & A4), and not in the period Before and After (edf = 1.32, F = 1.12, p = 0.237). The clicks per minute was lowest directly after echosounder activity, but reached stable levels about 5 minutes after the echosounder signal had stopped, after which they no longer significantly differed from the Before and After period, much faster than the ∼30 min recovery in acoustic presence of harbour porpoises The model explained 55.9% of the deviance (adjusted R^2^ = 0.515). The random effect of deployment also contributed significantly again to the model (edf = 11.84, F = 49.28, p < 0.001).

To gain insight into changing responses over the first 6 weeks of the During period, we compared harbour porpoise acoustic presence between weeks. The GAM again revealed a highly significant interaction between time of the hour and the week (edf = 27.54, F = 22.94, p < 0.001, Figure A6). The temporal pattern was again an increase in proportion PPM followed by stabilisation. However, pairwise comparisons did not show any difference in temporal patterns between weeks. A random effect smooth for deployment was significant again (edf = 11.94, F = 174.49, p < 0.001). The model explained 51.3% of the deviance (adjusted R^2^ = 0.494).

To investigate whether harbour porpoises returned to baseline acoustic presence levels towards the end of the 50-minute period in between echosounder activity, we compared the proportion of PPM in minutes 39–58 during the last week of echosounder activity with the first days after that period (7 days when available). A paired t-test showed no significant difference in PPM between the last week of echosounder activity and subsequent period after (t_6_ = −0.98, p = 0.363, mean difference = −0.0075, 95% CI [-0.026, 0.011], Figure 4). This suggests that harbour porpoise acoustic presence likely recovered to their typical levels during pauses in echosounder activity, or that effects are longer-lasting.

When examining patterns in harbour porpoise click types, to explore potential changes in foraging behaviour, we found – irrespective of time – no overall effect of period (estimate = 0.12 ± 0.11, p = 0.26). There were also no significant smooth effects over time bins of the hour, for neither periods (Before and After: edf = 1.53, F = 1.76, p = 0.284; During: edf: 1.00, F = 0.00, p = 0.956, Figure A7), indicating no consistent temporal pattern across the hour. The random effect of deployment also contributed significantly to the model (edf = 11.23, F = 15.06, p < 0.001). The model explained 51.9% of the deviance (adjusted R^2^ = 0.525).

## Discussion

By combining long-term monitoring of harbour porpoise calls with repeated on–off cycles of echosounder operation during 13 deployments, this study provides the first systematic assessment of the potential impact of high-frequency echosounders (for assessing pelagic fish abundance) on the vocal presence of harbour porpoises. We found the following answers to our questions: echosounder activity led to a significant reduction in the acoustic presence of harbour porpoises. There was a significant recovery within the pauses between subsequent echosounder activity events, reaching stable levels ∼30 minutes after the end of the echosounder activity. This stable level appeared to be the baseline level. The number of harbour porpoise clicks per positive minute also dropped, but recovered in ∼5 minutes, indicating that the drop in acoustic presence likely reflects porpoises leaving the area rather than solely a drop in click rate and thus poorer detectability. The ES70 transducer, operating at a nominal frequency of 70 kHz, appeared to have the strongest impact. Whether the ES200 transducer, with a frequency range of 185-255 kHz, alone would also cause a reduction in acoustic presence by itself cannot be derived from the current data, but seems unlikely given that recovery from ES70 exposure was already underway during ES200 operation, and as most of the ES200 signal lies above the upper hearing range of harbour porpoises. There is no sign of habituation over the weeks of hourly exposure events and we found no change in the relative amount of foraging buzzes. Lastly, we found no change in acoustic presence at 2.5 km distance from the echosounders. This means that the effect does not likely extend to this range, but our deterrence findings should be taken into account in specific studies on local assessments of predator-prey relationships. They should also raise awareness for a potentially accumulative impact from the wide use of echosounders in offshore practices on sensitive cetacean species.

### Harbour porpoise deterrence

We found a strong reduction in the acoustic presence of harbour porpoises during and immediately after echosounder activity, with subsequent recovery within the 50-minute periods in between transmissions. This pattern was consistently observed across all locations, and was validated across harbour porpoise detection devices (C/F-PODs and SoundTraps). The number of clicks detected in porpoise positive minutes also decreased, but less strongly and only briefly, suggesting that the found reduction in acoustic presence is either caused by displacement or complete cease in vocal activity rather than reductions in click rates and subsequent decreased detection capability.

Nevertheless, we cannot definitively conclude whether porpoises ceased vocalizing or left the area based on our data alone. It is well established that harbour porpoises typically produce clicks almost continuously at a relatively constant rate (Akamatsu et al., 2007; Osiecka et al., 2020; Sørensen et al., 2018a). Given the highly directional nature of these clicks (Wisniewska et al., 2015), movement away from the detection device (and echosounder) could reduce detectability (Macaulay et al., 2023) before the animals have fully left the area. There is some evidence that click rates can be affected by noise exposures. In a field study involving tagged wild harbour porpoises exposed to an acoustic deterrence device; one individual strongly increased its click rate, while four others reduced theirs – though still averaging 261 clicks per minute. Two of these individuals also showed decreased click amplitude (Elmegaard et al., 2023). Another field study, this time on the impact of vessel noise on seven tagged porpoises, reported a cease in regular echolocation clicks for one individual during a specific ferry pass, but systematic analyses focussed on foraging buzzes rather than other echolocation clicks (Wisniewska et al., 2018). Lastly, a study on two captive harbour porpoises found that click rates decreased in only 3 out of 25 exposures to pinger-like sound, typically during the first exposure to a novel sound type (Teilmann et al., 2006). Note that these studies examined click behaviour during noise exposure, whereas most of our observations are based on the quiet periods between echosounder activity. While the current reported reduction in acoustic presence may partly reflect changes in vocal behaviour, amplitude, or orientation, the consistently high baseline click rates reported in literature in combination with the only brief reduction in clicks per minute after the echosounder end, suggest that the observed reductions are primarily due to porpoises temporarily leaving the area. Studies with tagged porpoises can provide a definitive answer to this.

Previous studies revealed various behavioural responses of harbour porpoises to human-made sound (Erbe et al., 2025a). Ship noise, for instance, has been linked to changes in swimming and diving behaviour and interrupted foraging behaviour (Frankish et al., 2023; Wisniewska et al., 2018), whereas recreational boats only caused short and minor responses (Hao et al., 2024). Impulsive sound from seismic surveys and piling have led to changes in swimming behaviour including displacement (Benhemma-Le Gall et al., 2021; Haelters et al., 2015; Kastelein et al., 2019; Pirotta et al., 2014; Thompson et al., 2013). The impact of sonar is mostly studied in captivity and primarily focused on low and mid-frequency sonar (<10 kHz), to which porpoises typically responded with startle responses, increased swimming speed and respiration rate (Elmegaard et al., 2021; Erbe et al., 2025a; Kastelein et al., 2014). One example involving higher frequency sonar (25 kHz) showed that a captive harbour porpoise increased its respiration rate up to 150% depending on sound level and pulse type, and was reported to resume normal behaviour immediately after exposure (Kastelein et al., 2015). Other higher-frequency signals have been tested in the context of acoustic deterrence from fisheries or piling and reported displacement from ∼1.8 km (Hiley et al., 2021) up to ∼12 km (Dähne et al., 2017). We are only aware of one other study that looked into the impact of echosounders on harbour porpoises, as potential contributor of the impact of ship passes: Only echosounders of 200 kHz were detected, and they did not increase the likelihood of a response to the ship passes (Dyndo et al., 2015) (but see further ‘Echosounder types’). Studies on other cetaceans found no (Varghese et al., 2021) or subtle behavioural effects (Quick et al., 2017), and two studies on beaked whales found a decrease in acoustic presence (Cholewiak et al., 2017; Trickey et al., 2022), in one of the studies likely because of a cease in foraging as no significant drop in visual sightings was found (Cholewiak et al., 2017). To our knowledge, the current study is unique in its focus on a relatively high-frequency sonar and a very high-frequency hearing cetacean, revealing strong reductions in vocal presence, likely due to displacement.

### Recovery

During the 50-minute pauses between echosounder activity, the acoustic presence of harbour porpoises initially increased and stabilized after approximately 30 minutes. When comparing this stabilized porpoise presence in the final week with echosounder activity to the porpoise presence in the first week (3-7 days) without echosounder activity, we found no significant difference, suggesting that porpoise presence recovered to levels typical for the location and time of the year during the pauses.

Fast recovery from sound exposures has been reported before, especially in captive harbour porpoises. Several studies in captivity reported diminished responses already during 30-minute sound exposures (Kastelein et al., 2013), or an immediate return to the previously avoided area after the exposures (Teilmann et al., 2006) as well as a rapid return to baseline behaviours (Kastelein et al., 2013, 2000; Teilmann et al., 2006). However, in captive settings, porpoises are likely unable to fully avoid noisy conditions, and recovery (and habituation) patterns may therefore differ from those of free-ranging animals. In the field, porpoise detections have been reported to recover to baseline levels within 5 h after piling (Dähne et al., 2017), 24 h after piling (Brandt et al., 2011), and porpoises were detected again a few hours after seismic surveys (Thompson et al., 2013). In both the current and the mentioned field studies, it remains unclear whether the same individuals returned after initial displacement, or whether recovery just reflected (a normal rate of) new porpoises entering the area. These studies show that recovery might be fast, but also emphasize the need for individual-level tracking to disentangle return from replacement.

### Responsiveness over time

To examine whether porpoises reduced their responsiveness to echosounders over time – potentially as a consequence of habituation, desensitization, or motor fatigue – we compared their responsiveness over six weeks of echosounder use. We found no significant differences, indicating no changes in responsiveness over six weeks of hourly exposure. Note that these patterns are examined per location, and not individual-level patterns in responsiveness. Captive studies have shown that responses to noise can diminish over sequential exposures. For example, displacement effects to pinger-like sounds were strongest during the first few trials of a given sound type (Teilmann et al., 2006), and heart rate drops during sonar-like exposures also diminished over the first few trials (Elmegaard et al., 2021). Nevertheless, responses can also persist in captivity: four porpoises in a net pen responded to almost 30% of the passing boats, despite long-term residence in the harbour (Dyndo et al., 2015). Similarly, in a field study with tagged porpoises, the responses to loud boats persisted despite frequent exposure (Wisniewska et al., 2018). However, some reductions in responsiveness have also been documented in the wild, such as to a pinger (Cox et al., 2001), and during two 10-day seismic surveys (Thompson et al., 2013). Taken together, these findings suggest that some degree of habituation or reduced responsiveness over sequential exposures can be expected, but some responsiveness may persist. For free-ranging individuals like those in our study, habituation may be limited because animals can choose to temporarily leave the area – reducing their exposure and potential development of habituation – and naïve individuals may enter, maintaining a level of responsiveness in the local population.

### Echosounder types

We found that porpoise presence began to recover during the use of the ES200 transducer, and that recovery was slightly – but significantly – delayed at the two locations where the ES70 transducer was used instead of the ES200 (in minute 5-9). This suggests that the ES70 transducer has a stronger deterrent effect on harbour porpoises. Because the ES200 was always deployed following the ES70 in our study, we cannot determine whether the ES200 alone would also elicit a deterrent response. However, another study examining the response of four harbour porpoises in a net pen to passing ships, found no differences in responses to ships with and without a 200 kHz echosounder (Dyndo et al., 2015).

Behavioural audiograms indicate that harbour porpoises can detect frequencies up to at least 180 kHz, and are most sensitive in the 16 to 140 kHz range (Kastelein et al., 2002). This range includes the bandwidth used for communication and echolocation (110-150 kHz; Clausen et al., 2011; Miller, 2010). In this study, we set the ES200 transducer to emit frequencies starting at 185 kHz, and measured sound output from ∼192 kHz onwards (Figure 2). Additionally, the ES200 appears to have a stronger upward directionality (Simrad, n.d., n.d.) and was operated at a lower power setting than the ES70 transducer. Altogether, this may mean that harbour porpoises did not – or only poorly – detect the ES200 transducer. In contrast, the ES70 transducer operated well within the porpoise’s most sensitive hearing range, confirming it contributed more strongly – or solely – to their behavioural response.

Extrapolating effects of the specific echosounders examined in this study to echosounders in general must be done with caution (Hubert et al., 2024), but a preliminary risk assessment of the potential impact of echosounders on marine mammals can likely be made by considering several key factors: the echosounder’s frequency range and overlap with the hearing sensitivity of marine mammals present in the area, its amplitude, beam width, and the attenuation outside the main beam (Figure 5). Most fish-finding echosounders operate at relatively high frequencies (38-200 kHz; MacLennan and Simmonds, 2013), partially overlapping with the most sensitive hearing ranges of high-frequency (best hearing up to ∼110 kHz) and very high-frequency (best hearing up to ∼140 kHz) cetacean hearing groups; which includes all studied toothed whales (Southall et al., 2019). In contrast, low-frequency hearing cetaceans (baleen whales) and other marine mammals primarily hear lower frequencies with best hearing up to 19 kHz and 8.3-30 kHz respectively (Southall et al., 2019), and therefore show less overlap with most typical echosounder frequencies. Furthermore, echosounders that produce moderate sound levels with narrow beams and steep attenuation outside the main lobe are likely to exponentially reduce the number of individuals exposed to potentially disturbing sound levels. Although the overall prevalence of sonar use remains unknown, it is expected to be greatest among naval, fishing, research, and commercial survey vessels. In a dataset of close to 16,000 commercial vessel transits near the Port of Vancouver (Canada), 20–60 kHz sonar was detected in 1.3% of cases, with higher frequencies undetectable due to sampling limitations (Martin et al., 2021).

**Figure 5:**
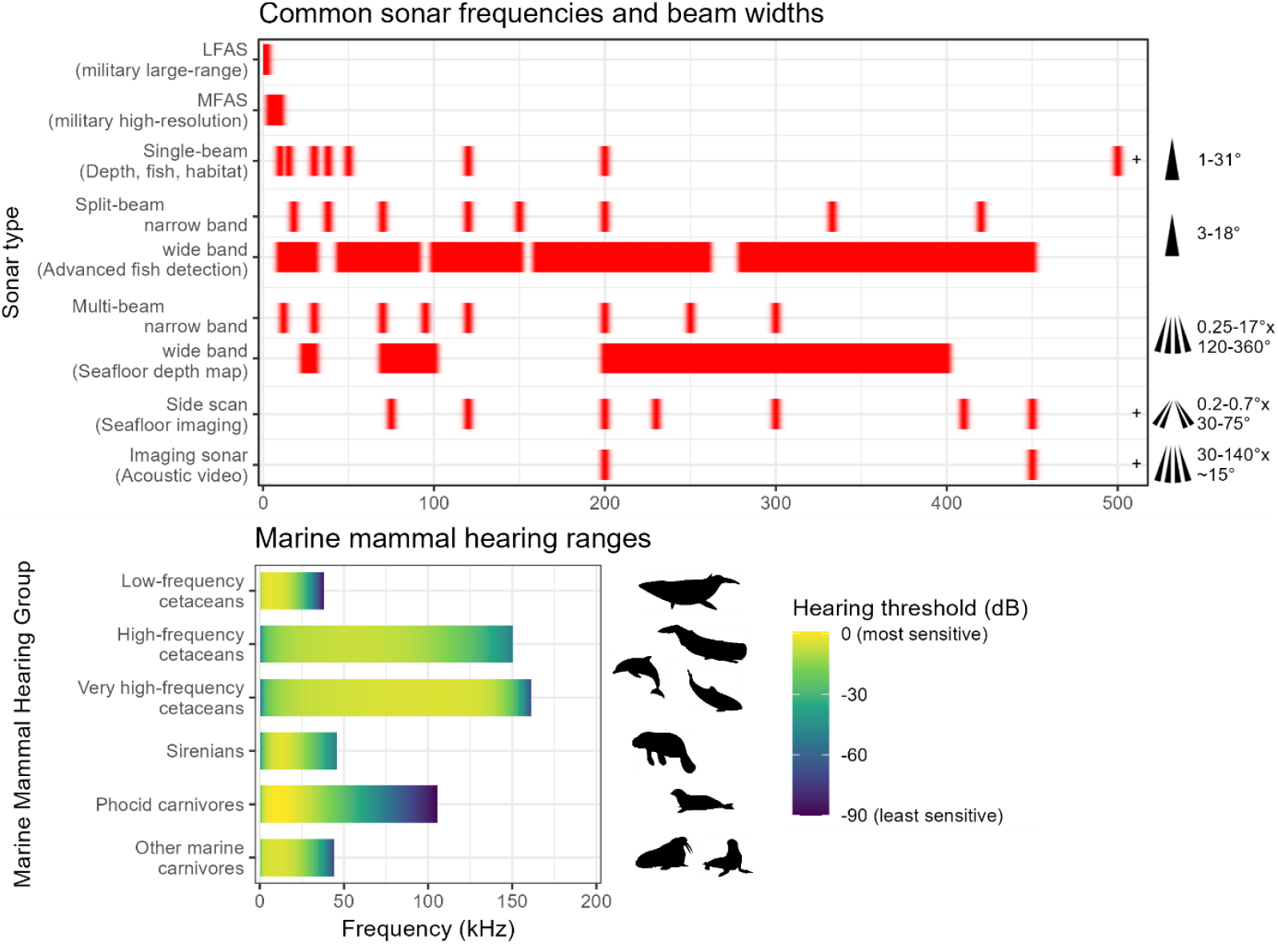
Overview of the common frequency ranges and beam widths of common sonars, including echosounders (top panel), and hearing ranges of marine mammals (bottom panel). Frequency ranges and beam widths of sonars are based on an inventory of systems currently for sale. For single-beam echosounders, side scan, and especially imaging sonars, even higher-frequency systems are available (indicated by the +). Hearing ranges and relative sensitivities are based on Southall et al., 2019. All marine mammals can detect military sonar and other relatively low-frequency systems. Somewhat higher-frequency systems can also be detected by (very) high-frequency hearing cetaceans and phocids, although with decreasing sensitivity for the latter group.

### No changes in foraging

We found no changes in the proportion of buzz-clicks over time during the 50-minute pauses, nor any differences between the period with and without echosounder activity, indicating no lasting changes in foraging behaviour. Minutes during active echosounder use were excluded from the analysis due to limited porpoise detection capability during use of the ES70 transducer, and very few detections during use of the ES200 transducer. It is important to note, however, that buzz-click proportions outside these periods reflect individuals that either stayed or returned to within the detection range of the C/F-PODs, and may therefore be biased toward less-affected individuals. Nevertheless, Pirotta and colleagues (2014) also found a decrease in buzz-clicks in harbour porpoises that stayed in a sound-affected area by seismic surveys. Another field study found decreases in buzz-clicks in four out of six tagged individuals during high levels of ship noise (Wisniewska et al., 2018). Given that porpoises must forage almost continuously to meet their high metabolic demand, their capacity to cope with repeated human disturbances is limited (Wisniewska et al., 2016). While we cannot exclude brief cessations in foraging, our findings indicate no persistent effect of echosounder use on buzz-click activity.

## Conclusion

This study demonstrates that bottom-moored echosounders can temporarily reduce the acoustic presence of harbour porpoises, likely indicating short-term deterrence. However, porpoise numbers consistently recovered approximately 30 minutes after the echosounder stopped. Despite fast recovery, no evidence of habituation or reduced responsiveness was found over a six-week period, but note that responsiveness is assessed per location, and not per individual. Among the two transducers tested, the ES70 (70 kHz) had a stronger deterrent effect than the ES200 (185-255 kHz), likely primarily due to its overlap with the porpoise hearing range and potentially also due to higher amplitude, beam width, and lower attenuation outside the beam. We found no indication of lasting changes in the porpoises’ proportion of buzz clicks during pauses between echosounder transmissions, suggesting their foraging behaviour was either unaffected or recovered quickly. These results show that echosounders used for fish detection have the potential to disturb cetaceans, yet this received relatively little attention to date. Our findings underline the importance of considering frequency and beam width when assessing impacts on (very) high-frequency hearing cetaceans, provide a foundation for future impact assessments and mitigation strategies when high frequency sonars are used, and should be taken into account for studies on predator-prey relationship involving echosounders.

## Acknowledgements

We thank the crew of the vessels RV Simon Stevin, Anteos, and Aquaflight for helping us to deploy and retrieve the moorings, and the personnel of the windfarms C-Power, Belwind, and Borssele for permitting us to enter the offshore windfarms and placing the moorings. This study is part of the APELAFICO project which is funded by NWO (NWA.1236.18.004) and Rijkswaterstaat (through HVP funding from the EU). This project makes use of data and infrastructure provided by VLIZ and funded by the Research Foundation—Flanders (FWO) as part of the Belgian contribution to the LifeWatch ESFRI (I002021N-LIFEWATCH). Lastly, we thank the Harbour Porpoise Network Borssele project for making their data available, the project is part of the Wozep offshore wind ecological programme of Rijkswaterstaat on behalf of the Dutch Ministry of Climate Policy and Green Growth.

