## Appendix for "Echosounders for fish detection disturb harbour porpoises"

Table A1: Summary of all the included multi-sensor mooring deployments, including their location, deployment period, and instruments and settings used.

| | | | WBAT | | SoundTrap | | |
|---|---|---|---|---|---|---|---|
| Location[1] | Period | Duration (days[2]) | Duration (days[2]) | Transducers[3] | Model | HF clicks used[4] | POD type |
| Belwind OWF, BE | Summer '21 | 71 | 48 | 70 & 200 | 300 HF | Yes | C-POD |
| | Summer '23 | 70 | 60 | 70 & 200 | 300 HF | - | C-POD |
| CPower OWF, BE | Summer '23 | 70 | 60 | 70 | 300 HF | Yes | C-POD |
| | Autumn '23 | 71 | 60 | 70 & 200 | 600 HF | - | C-POD |
| Borssele OWF, NL | Spring '23 | 40[6] | 40 | 70 & 200 | 4300 HF | - | C-POD |
| | Spring '23 | 40[6] | 40 | 70 | 300 HF | Yes | C-POD |
| Grafton SW, BE | Summer '21 | 71 | 48 | 70 & 200 | - | - | C-POD |
| | Summer '23 | 68 | 60 | 70 & 200 | 300 HF | - | C-POD |
| | Autumn '23 | 71 | 60 | 70 & 200 | 600 HF | Yes | F-POD |
| Birkenfels SW, BE | Summer '21 | 71 | 48 | 70 & 200 | 300 HF | - | C-POD |
| Gardencity SW, BE | Summer '23 | 71 | 60 | 70 & 200 | 4300 HF | - | C-POD |
| NCN 2404[5] SW, NL | Spring '23 | 40[6] | 40 | 70 & 200 | 300 HF | - | C-POD |
| NCN 189[5] SW, NL | Spring '23 | 40[6] | 40 | 70 & 200 | 300 HF | - | C-POD |

[1]: OWF = offshore windfarm, SW = shipwreck, BE = Belgium, NL = the Netherlands
[2]: Rounded to full days
[3]: Nominal frequency in kHz, for full descriptions and characterization, see the Material & Methods
[4]: Only when data was available for 60 min per hour
[5]: Unidentified shipwreck
[6]: The deployments are not included in the 'Before and After' period as the deployment lasted less than a day longer than the 'During' (echosounder active) period

Table A2: Overview of echosounder settings.

| Transducer | Beam angle (degree) | Frequency (kHz) | | Beam type | Pulse type | Transmit power (W) | Pulse length (us) | Ramping | Ping interval (s) |
|---|---|---|---|---|---|---|---|---|---|
| | | Nominal | Range | | | | | | |
| ES70-18CD | 18 | 70[1] | | Split beam | Continuous Wave (CW) | 150 | 256 | Fast | 0.3 |
| ES200-7CDK | 7 | | 185-255[1] | Split beam | Frequency Modulated (FM) | 75 | 1024 | Slow | 0.6 |

[1]: According to the software of the manufacturer. See PSD (Figure 2) for the full spectral information measured on the moorings.

Table A3: Summary statistics of the proportion porpoise positive minutes in the during period.

| Summary statistics, weighted for 13 deployments | Proportion PPM | | |
|---|---|---|---|
| | 2nd min after 10 min echosounder period | 2nd min after ES70 | min 39-58 of each hour |
| Mean | 0.0127 | 0.0076 | 0.0359 |
| Median | 0.0069 | 0.0021 | 0.0338 |
| 25th percentile (Q1) | 0.0021 | 0.0007 | 0.0186 |
| 75th percentile (Q3) | 0.0228 | 0.0090 | 0.0476 |

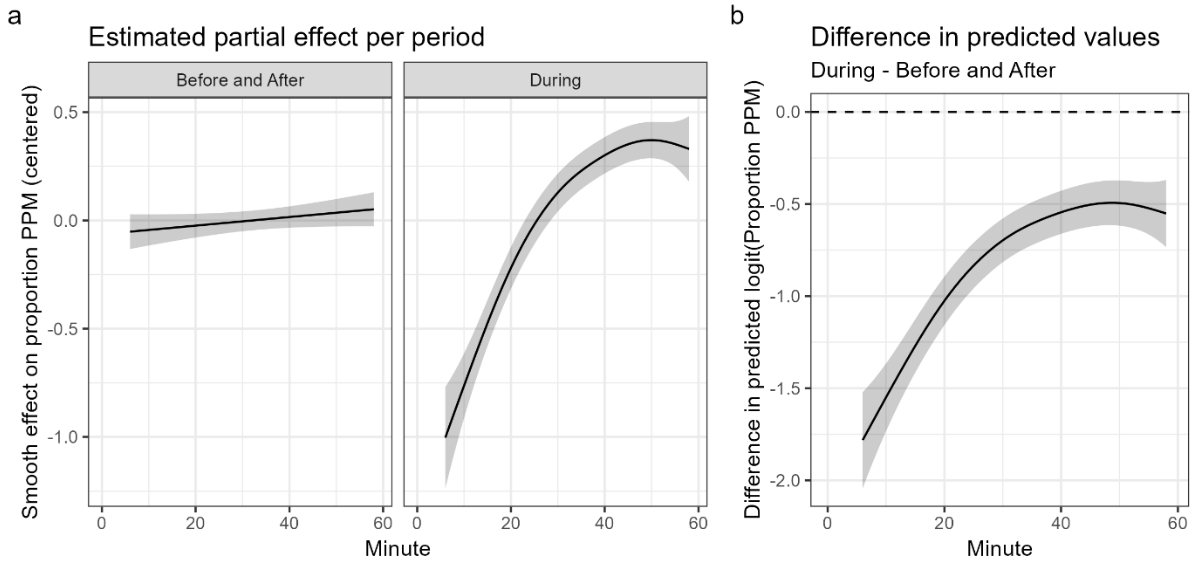


Figure A1: (a) Centred estimated partial effect on the relative proportion PPM over the hour, by period, based on the GAM. The solid black lines indicate the estimated smooth effects per period, and the shaded area indicate the 95% confidence intervals. For the before and after period, there was no significant within-hour trend in harbour porpoise presence. In contrast, for the During period (with echosounder activity), porpoise presence increases after the echosounder activity, and stabilizes in the final 20 minutes of the hour. (b) Estimated difference between the periods (During minus Before and After). The line and shaded bands show the difference on the link (logit) scale with 95% confidence intervals. The difference and 95% CI is below zero indicating a lower proportion PPM in the during period.

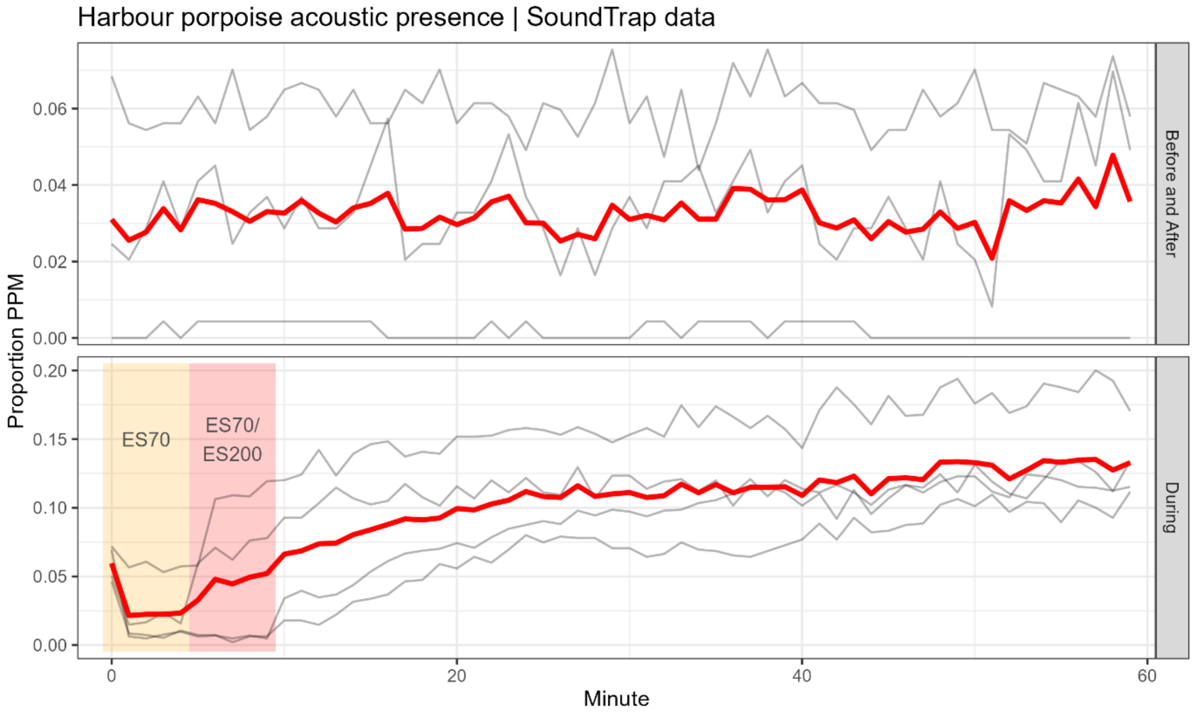


Figure A2: Proportion of harbour porpoise positive minutes (PPM) for each minute of the hour, based on SoundTrap high-click detections. Grey lines indicate individual deployments, and the solid red line shows the mean. We could use data from four deployments for the during period (Table A1), and three deployments for the before and after period, as cross-device validation of the pattern found using the C/F-PODs. Differences in absolute PPM between results from C/F-PODs and SoundTraps can likely be explained by differences in sensitivity and inclusion criteria.

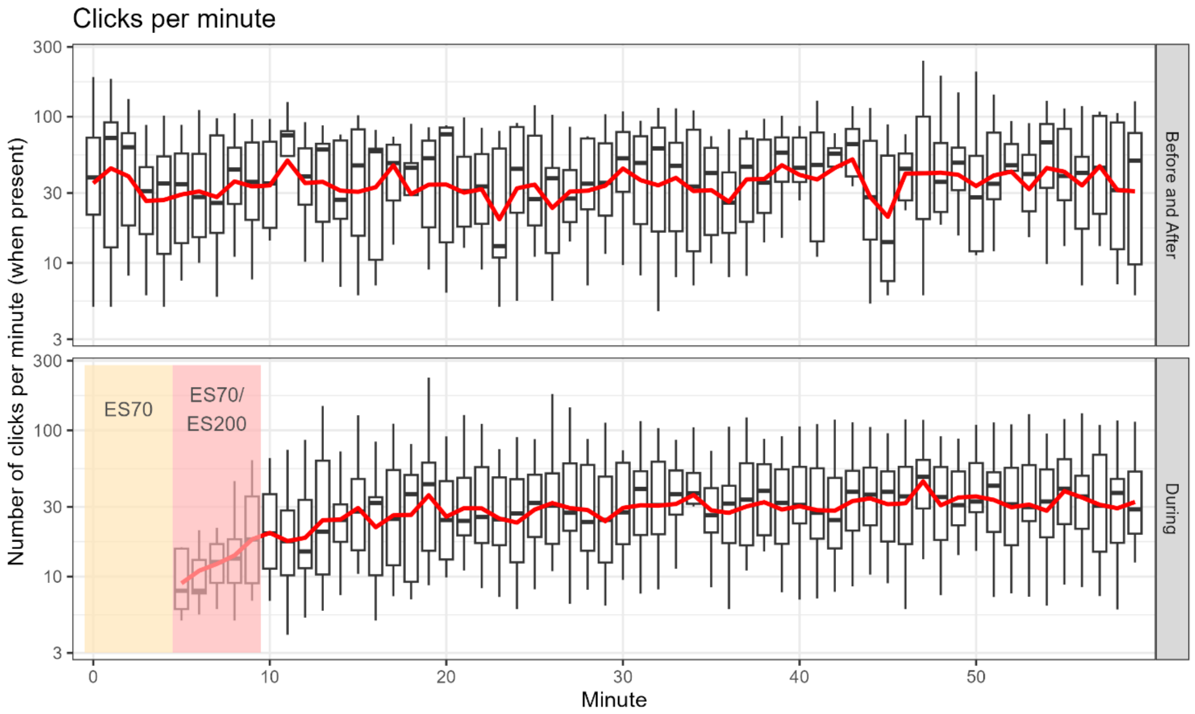


Figure A3: Mean porpoise click count per minute in which at one click is detected. Boxplots show the median, 25th and 75th quantile, minimum and maximum without outliers, and outliers of all locations. The solid red line indicates the weighted mean of all locations. Note that the y-axis has a logarithmic scale.

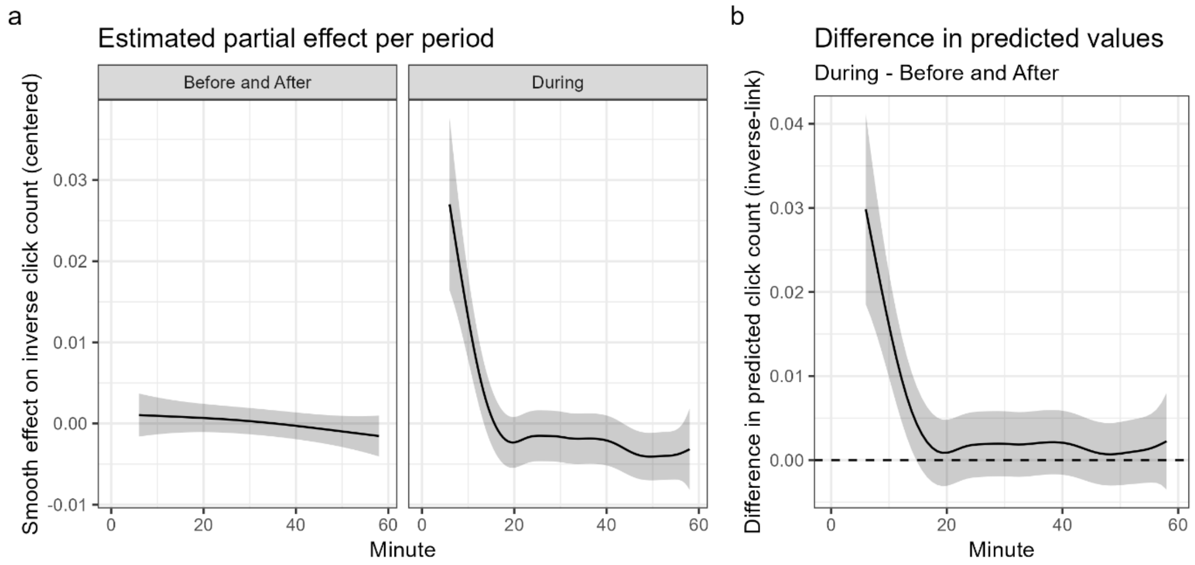


Figure A4: Estimated partial effect of time (minute of the hour) on the clicks per positive minute, by period, based on a GAM. The solid lines indicate the estimated smooth effects per period, and the shaded area indicate the 95% confidence intervals. In the during period, the number of clicks per minute reaches a stable – and the mean – level about 5 minutes after the end of the echosounder (b) Estimated difference between the periods. The line and shaded bands show the difference on the inverse-link scale with 95% confidence intervals. The first difference between click rate in the before and after period with the during period lasts until 4.9 minutes after the echosounder end.

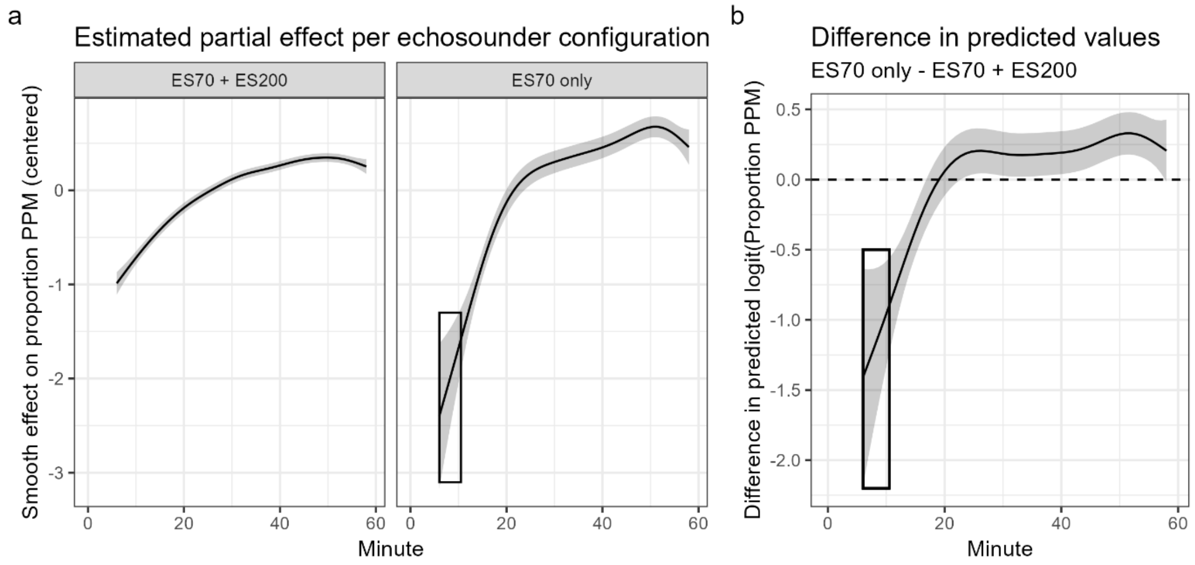


Figure A5: (a) Estimated partial effect of time (minute of the hour) on the relative proportion PPM, by echosounder transducer configuration, based on a GAM. The solid lines indicate the estimated smooth effects per transducer configuration, and the shaded area indicate the 95% confidence intervals. Similarly to the previous model, porpoise acoustic presence initially increases, followed by a stabilization. For the two deployments with the ES70 transducer only, the increase lags behind at first, but quickly catches up and ultimately even has a higher porpoise acoustic presence. The model output shown in the black square is outside the range of observed data and should be ignored. (b) Estimated difference between the configurations. The line and shaded bands show the difference on the link (logit) scale with 95% confidence intervals. The porpoise acoustic presence first stays behind in ES70 only deployments, but already caught up within 10 minutes and even reaches higher porpoise levels.

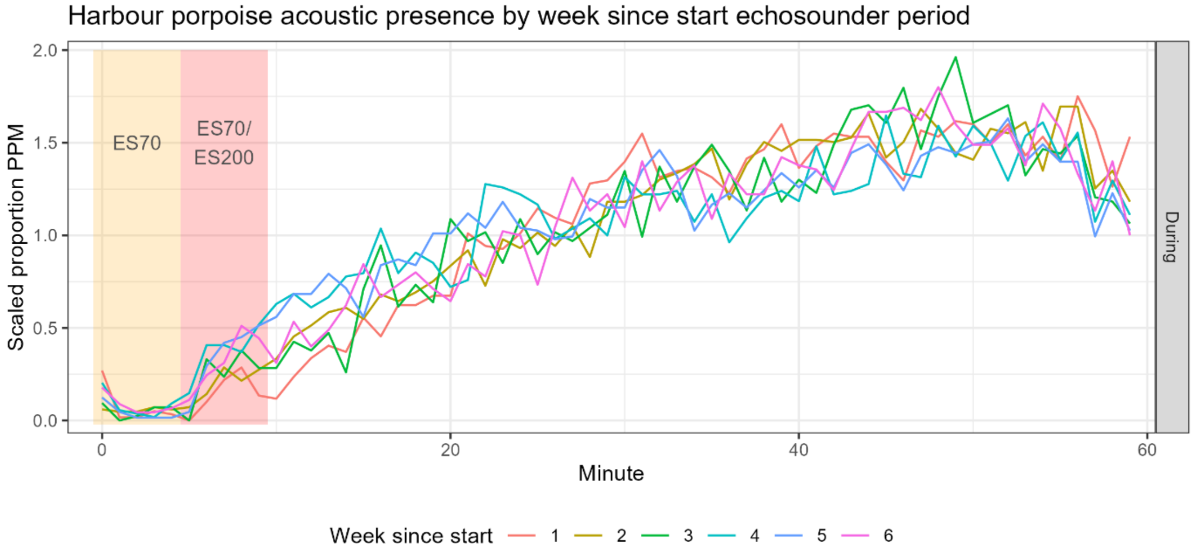


Figure A6: Scaled proportion of harbour porpoise positive minutes (PPM) within the hour and grouped by week since the start of the echosounder activity, only considering the period where the echosounder was active. Each line shows the scaled mean of 13 deployments. The shaded area indicates the minutes with echosounder activity.

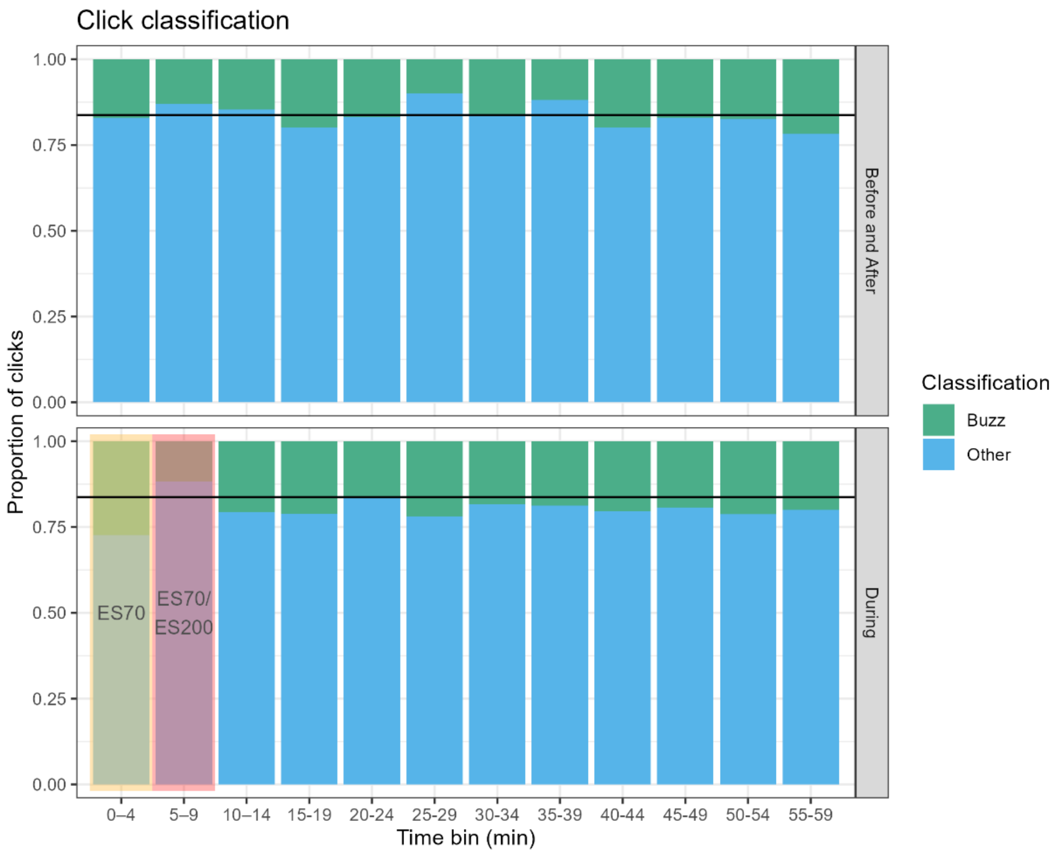


Figure A7: Classification of harbour porpoise click type (based on click interval). The shown proportions are the weighted means for all deployments, grouped in 5-minute bins. The horizontal black line shows the mean proportion of 'other' clicks in the 'before and after' period. The shaded area indicates the minutes with echosounder activity. During minutes with an active ES70 transducer, click detections are incomplete because of C-POD saturation.
